# Understanding the mechanisms underlying microbiota variation in wild tick populations

**DOI:** 10.1101/2024.12.27.630517

**Authors:** Víctor Noguerales, José de la Fuente, Sandra Díaz-Sánchez

**Affiliations:** Departamento de Biología Animal, Edafología y Geología, Universidad de La Laguna, San Cristóbal de La Laguna, Tenerife, Spain; Instituto de Productos Naturales y Agrobiología (IPNA-CSIC), San Cristóbal de La Laguna, Tenerife, Spain; SaBio. Instituto de Investigación en Recursos Cinegéticos IREC(CSIC-UCLM-JCCM), Ronda de Toledo s/n, 13005 Ciudad Real, Spain; Department of Veterinary Pathobiology, Center for Veterinary Health Sciences, Oklahoma State University, Stillwater, OK 74078, USA

**Keywords:** 16S metagenomics, Ixodidae, microbial ecology, microbiota, ticks

## Abstract

Ticks are fascinating arthropods that have developed a unique blood-feeding lifestyle, making them intriguing subjects for understanding how they adapt to living with vertebrate hosts, their ability to transmit diseases, and their versatile roles in the ecosystem. Recent research highlights how significantly microbes impact the evolution of ticks, suggesting a genetic link to their hosts. These microbial partnerships are vital in improving tick nutrition, metabolism, reproduction, and overall survival, highlighting their important role as vectors in diverse ecological environments. In this study, we took a close look at the microbial communities present in three wild tick species—*Hyalomma lusitanicum*, *Rhipicephalus sanguineus*, and *Rhipicephalus bursa*—collected from several sites across central Spain. Through high-throughput DNA sequencing of the 16S rRNA gene, we discovered a complex and diverse microbiota largely composed of Proteobacteria, Bacteroidota, and Firmicutes. While all species exhibited a shared core microbiota, notable differences in microbial composition and network microbial interactions suggest that each species may be influenced by ecological and evolutionary factors shaping their microbiota. These insights deepen our understanding of the complex relationships between ticks, their microbiota, and the surrounding environment, which can inform better strategies for managing vectors and controlling pathogens.

## Introduction

Ticks (Acari: Ixodida) are arachnids that transitioned to a blood-feeding lifestyle approximately 250 million years ago (Mans and Neitz, 2004). This evolutionary path has turned ticks into excellent model organisms for examining their adaptations to vertebrate hosts, the development of vector abilities, and their ecological versatility (Jia et al., 2020; Mans et al., 2023). In the early 21st century, Zilber-Rosenberg and Rosenberg (2008) emphasized the essential role of microbes in host evolution, suggesting that microbes should be considered a genetic component of the host. This well-established idea highlights the importance of symbionts and endosymbionts in the evolutionary adaptability of ticks as vectors. These microbial alliances aid in tick nutrition (e.g., supplying vitamins and cofactors, enhancing blood-feeding), metabolism (e.g., facilitating nitrogen recycling, respiration, and osmoregulation), reproduction (e.g., boosting success and larval motility), and survival (e.g., enhancing temperature tolerance) (Neelakanta et al., 2010; Ahantarig et al., 2013; Vavre et al., 2014; Narasimham et al., 2014;2015; Kolo et al., 2023).

Research on tick-microbe interactions has advanced our understanding of tick biology and vector competence. These interactions reveal dynamics at the microbiome-pathogen interface, impacting pathogen transmission. For example, Narasimhan et al. (2014) showed that disrupting the gut microbiota of *Ixodes scapularis* larvae enabled colonization by *Borrelia burgdorferi*, the cause of Lyme disease. Importantly, *Borrelia*-positive ticks from natural populations displayed greater bacterial diversity compared to *Borrelia*-negative ticks. These findings highlight the importance of microbial diversity in pathogen dynamics and vector competence. Another example of interactions between ticks, microbiota, and pathogens is seen with the obligate intracellular pathogen *Anaplasma phagocytophilum,* which alters tick proteins to disrupt gut microbiota, thereby facilitating its transmission to the salivary glands (Abraham et al., 2017). Beyond facilitating pathogen transmission, tick-microbiota interactions may drive novel adaptations, reproductive isolation, and speciation (Gilbert et al., 2015; Mazel et al., 2018). For example, Neelakanta et al. (2010) found that *A. phagocytophilum* infection increased tick tolerance to freezing by inducing antifreeze glycoproteins production.

The composition of tick microbiota is influenced by various factors, including life stage, feeding status, and local environmental conditions that differ by geographical context (Bonnet & Pollet, 2021; Aivelo et al., 2022). However, the specific processes that shape and organize a tick’s microbial community remain unclear. Ticks may obtain their microbiota through maternal transmission and horizontal transfer from their surroundings, which could occur through the spiracles, mouth, and anal pore, or even during copulation (a paternal transmission route) or while feeding on host blood (Narasimhan & Fikrig, 2015; Bonnet & Pollet, 2021; Pavanello et al., 2023). The local environment serves as a significant microbial reservoir, fostering interactions that stimulate arthropod development and microbial assembly (Guégan et al., 2018; Hannula et al., 2019; Girard et al., 2023). Although not extensively studied, elements from tick hosts and their microbial interactions may also affect tick microbiota composition (Frazenburg et al., 2013; Davenport, 2016). Only a limited number of studies identify species-specific connections in tick-related microbiota, particularly regarding a close evolutionary relationship between ticks and certain microbes (Cabezas-Cruz et al., 2018; Díaz-Sánchez et al., 2019). Exploring species-specific microbiota within wild tick populations might offer valuable insights into the factors influencing microbial diversity, crucial for understanding vector competence, pathogen transmission, nutritional support, reproductive success, environmental stress resilience, and immune regulation (Bonnet et al., 2017; Bonnet & Pollet, 2021; Narasimhan et al., 2021).

This study employed high-throughput DNA sequencing to examine the microbial composition of wild tick species, recognizing potential variations among sympatric species. Overall, the findings indicated a rich and diverse bacterial microbiota associated with wild ticks, implying the existence of a stable core group of bacteria. Nonetheless, notable differences between tick species may arise, underscoring the specificity of the microbiota and supporting the idea that microbial communities are influenced by a mix of abiotic factors (such as climate and vegetation) and biotic interactions, including competition among microbes and host-specific dynamics.

## 2. Material and methods

### 2.1 Tick sampling and identification

We used the flagging method to capture ticks in five different locations in Castilla-La Mancha (Spain) during the spring-summer-autumn season (2019-2020). We selected sites where diverse species of ticks could be found to maximize sympatry among the species included in the study (**Table 1**). All ticks collected were taxonomically identified at the genus and species levels using a stereomicroscope and following the taxonomic keys described by Manilla 1998. They were confirmed through individual genetic analyses via Sanger sequencing of the mitochondrial COI gene, as well as the 16S rRNA and 12S rRNA genes (**Table 2**). To prevent environmental microbial contamination, all collected ticks were placed individually in new, sterilized 1.5 mL microcentrifuge tubes following a surface sterilization protocol in the laboratory involving successive washes: brief washes of 1 mL 3% hydrogen peroxide (1 minute vortex), 70% ethanol twice (30 seconds each), and phosphate buffered saline solution (2 minutes). Following surface sterilization, each tick was placed in a new, sterilized 1.5 mL microcentrifuge tube and kept frozen at −20 °C before DNA extraction.

**Table 1.**
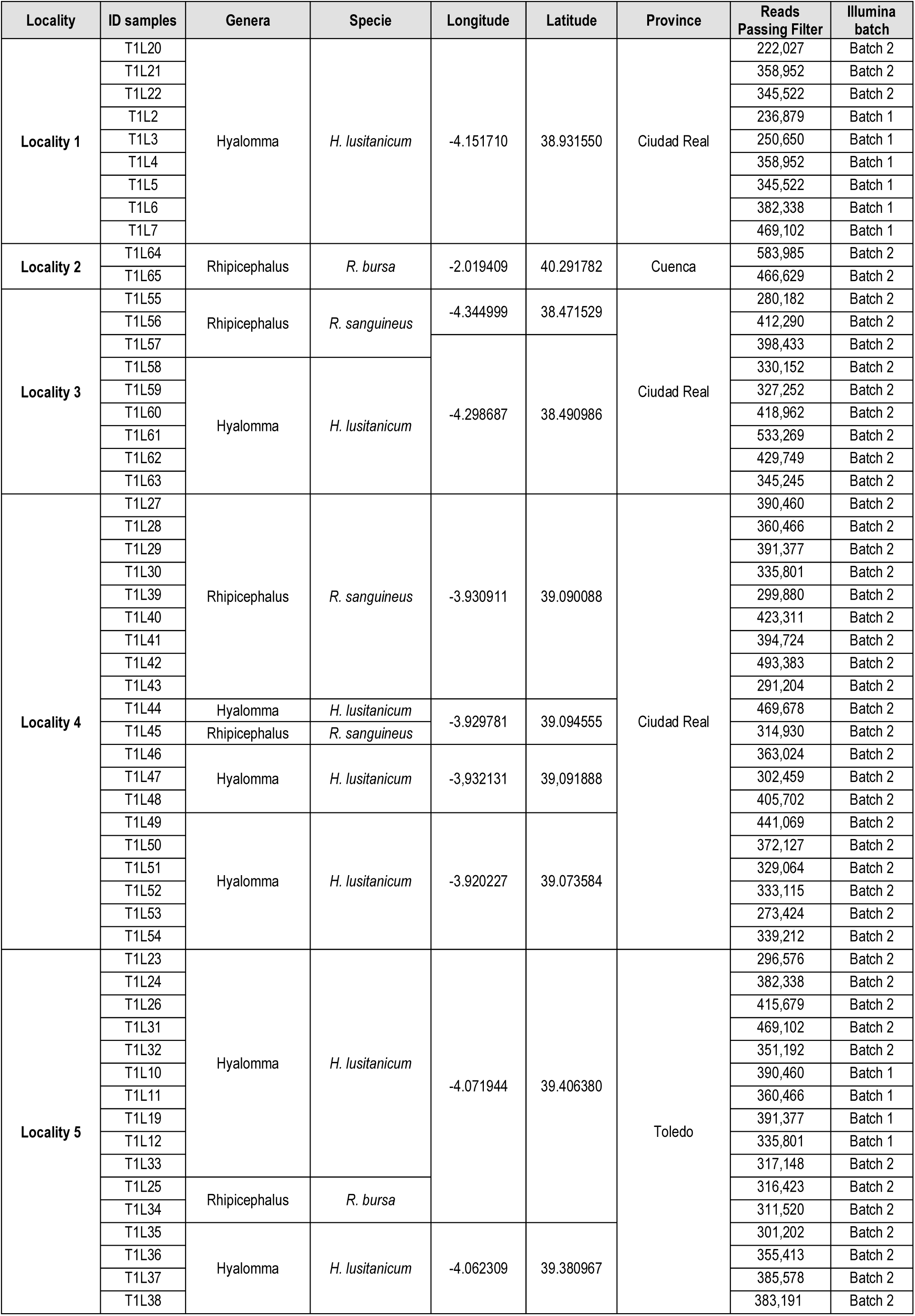
Description of samples, collection sites and sequence performance.

**Table 2.**
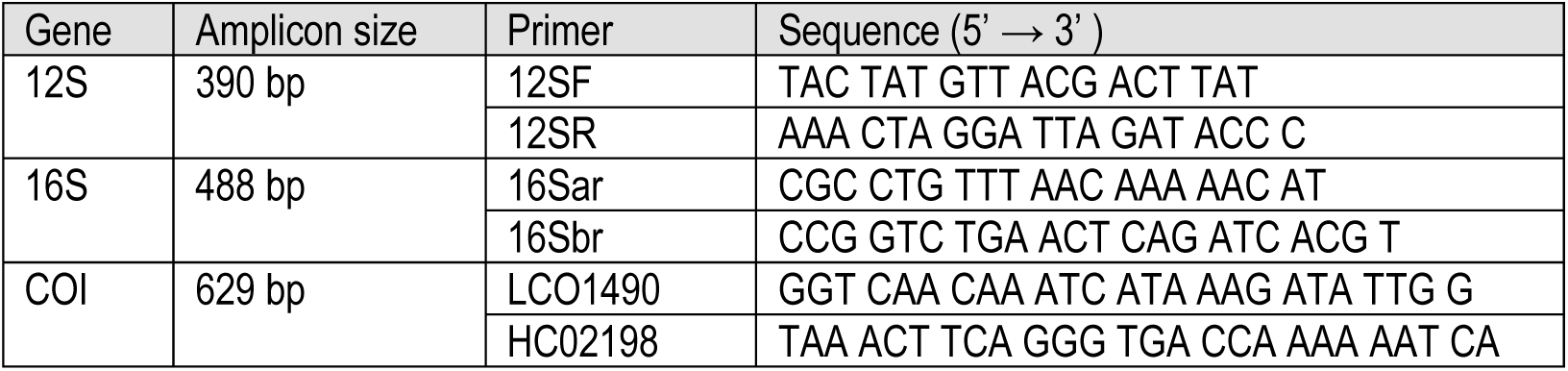
mtDNA genes, PCR amplicon sizes, primers used for amplification, and their respective sequences (Simon et al. 1994).

### 2.2 DNA extraction, library amplicon preparation, and sequencing

Following purification, DNA samples were sent to the Genomic Unit at Campus Moncloa (University Complutense of Madrid) for sequencing analysis. An aliquot from each DNA sample was used to prepare libraries that amplified the V3 and V4 hypervariable regions of the 16S rRNA gene, employing the primer pairs published by Klindworth et al. (2013). PCR amplification of the target amplicon was carried out according to the manufacturer’s guidelines. The anticipated PCR product size, approximately 480 bp, was verified using a Bioanalyzer DNA 1000 chip (Agilent, US) and further purified with AMPure XP beads (Beckman Coulter, Life Sciences, US) for subsequent processing. Next, Illumina sequencing adapters and index barcodes were added to the amplicon target using the Nextera XT DNA Library Preparation Kit (Illumina Inc, CA, US), after which the libraries were pooled for sequencing. All cluster generation and paired-end sequencing were executed on the Illumina Next Generation sequencing MiSeq system, following the Illumina MiSeq v2 2×300 cycle chemistry protocols. In total, we sequenced samples from 58 individuals across two sets: those collected in 2019 (batch 1=10 individuals) and 2020 (batch 2=46 individuals) (**Table 1**).

### 2.3 Sequence and microbiota analysis

A total of 17,062,421 Illumina MiSeq reads in R1 and R2 passing filter (**batch 1**= 487,529; average=48,752.9 per/sample; **batch 2=** 17.062.421; average= 370.922) were paired-end demultiplexed, and a fastq file was generated using Illumina MiSeq Reporter software (**Table 1**). Quality assessment was performed using the CLC Genomics Workbench (Qiagen). The raw 16S rRNA sequences were uploaded to the SRA repository (accession No. needs to be submitted). Sequence analysis was performed using the DADA2 (v1.12) inference algorithm on primer-free reads to correct sequencing errors and create amplicon sequence variants (ASVs) for the tick microbial communities in R (v4.0.1) (Callahan et al., 2016). The reads were quality-filtered using the *FilterAndTrim* function, which truncated the forward and reverse reads at 280 bp and 255 bp for the tick microbiota dataset. Further, reads were removed if they had more than 2 errors in the forward and 2 errors in the reverse reads. Reads were merged after the inference of sequence variation with *learnErrors* and denoised using DADA2 functions. Chimeric sequences were eliminated with *RemoveBimeraDenovo,* and taxonomy was assigned to ASVs using the classify-learn naïve Bayes taxonomic classifier *assignTaxonomy* based on the SILVA database (v138) (Yarza et al., 2014). Count abundances of taxa were extracted from original outputs for each taxonomic level (**Supplementary file 1**). Microbial community profiles were constructed at the phylum, family, and genus levels for further analysis. Because we detected a significant effect of sequencing batch on our diversity estimates, we performed all subsequent analyses on the subset of the data that included only the second batch of samples (referred to as “batch 2” in **Table 1**).

### 2.4 Statistical analysis of tick microbial communities

For the tick microbiota dataset, the ASV count table was generated. A total of 24,521 ASVs were assigned for a total of 46 samples (**Supplementary file 1: Table 1**) and at Phylum, Family and Genus taxonomic. All the subsequent biological analyses were performed using *phyloseq* (v.1.36) (McMurdie and Holmes, 2014) and the *ggplot2* package (Wickham, 2016) for visualizations in R (v4.0.1). At this point, we performed technical filtering to remove spurious ASVs in the bacterial dataset, which involved removing phyla with only 1 or 2 ASVs represented, followed by relative abundance filtering using a threshold of 0.05%. To achieve a high-quality representation of tick microbiota, we proceeded with several additional filtering steps. We excluded ASVs that do not appear more than 5 times in more than 2.5 samples in our dataset. Finally, we generated a *phyloseq* object with an *asv_table* of 1,356 taxa for downstream analysis. Then, the microbial community composition was represented in terms of relative abundance at the phylum, family, and genus levels, keeping the five most abundant taxa featured at each level using the tax-glom function in the *phyloseq* package. Core and shared genera between tick species and within localities where ticks were found in sympatry (locality 3, locality 4, and locality 5) were tallied using the *amp_venn()* function of the ampvis2 package (v2.7.28) (Andersen et al., 2018), considering the “*core*” genera shared by groups with an abundance ≥ 0.05% and present in 85% of the samples.

### 2.5 Tick microbiota diversity

We assessed the within-individual diversity (alpha-diversity) of tick species (*H. lusitanicum, R. sanguineus*, and *R. bursa*) and genera (*Hyalomma* spp, *Rhipicephalus* spp) using the Shannon-Wiener index, Gini-Simpson index (1-Simpsońs original index), and InvSimpson index through the *estimate_richness*() function in *phyloseq* (v.1.36) (McMurdie and Holmes, 2014). The Shannon index evaluates both ASV richness (number of ASVs) and evenness (distribution of ASV abundances), indicating the uncertainty about the identity in a sample. Lower diversity implies lower prediction uncertainty, as a randomly selected species is likely to be the dominant one. In contrast, higher diversity correlates with increased uncertainty. The Gini-Simpson index assesses the chance that two randomly selected members from the same sample belong to different species, with scores ranging from 0 to 1; a lower score suggests reduced diversity. The InvSimpson index complements the Gini-Simpson and ranges from 1 upwards, with higher values indicating greater diversity. Non-phylogenetic diversity metrics were visualized using the *ggplot2* package (v 3.5.1) in R, utilizing *geom_violin*(). Prior to estimating alpha diversity, we rarefied the dataset through random subsampling with *rarefy_even_depth*() to prevent low and uneven depth sequences (Hughes & Hellmann, 2005; McMurdie & Holmes 2013). Additionally, we included phylogenetic diversity metrics utilizing the *phyloseq_phylo_div*() function found in the *metagMisc* package for R (v 4.4.1). Given that phylogenetic diversity measures are not statistically independent of species richness (Tucker and Cadotte, 2013), we computed the standardized effect size of phylogenetic diversity, ses.PDFaith (referenced as Faith’s PD) (Faith 1992), along with the standardized effect size of mean pairwise distance within communities, ses.MPD (Mean Pairwise Distance among all species in an assemblage) (Webb et al. 2000, 2002). The standardized effect size metrics indicate that positive values suggest overdispersion, zero indicates random phylogenetic structure, and negative values point to phylogenetic clustering. Thus, ses.PD and ses.MPD evaluate the relative excess (overdispersion) or deficit (clustering) of phylogenetic diversity for a specific set of species in relation to the previously presented tree. ses.MPD is influenced more by the basal structure of phylogenetic assemblages across various evolutionary scales, whereas ses.PD better reflects the terminal structure of the tree. Utilizing both metrics together improves our comprehension of assemblage structure, especially when different processes operate concurrently at various scales (e.g., overdispersion, clustering, overdispersion of clusters, and clusterization). We visualized the phylogenetic diversity metrics using *ggviolin*() from the *ggpubr* package (v 0.6.0) R.

Subsequently, we conducted tests to identify statistical differences in the alpha-diversity metrics among tick species (*H. lusitanicum, R. sanguineus*, and *R. bursa*) by utilizing the non-parametric Kruskal rank test, employing the *kruskal.test* () function provided in the *ape* package (v 5.5; Paradis and Schliep, 2019). Additionally, we performed a post-hoc analysis for multiple comparisons through the *dunn.test*()function available in the *dunn.test* package (v 1.3.5; Dinno, 2017), adjusting the p-values in accordance with the Benjamini-Hochberg method. Furthermore, we statistically examined the presence of differences in the alpha-diversity metrics between the tick genera (*Hyalomma* spp. vs *Rhipicephalus* spp.) using the Wilcoxon rank sum test with continuity correction, implemented via the *wilcox.test*() function within the *stats* package (version 4.4.1) in R.

To determine whether the structure of tick microbiomes varies among species and genera (specifically the impact of host species on community dissimilarity), we assessed beta-diversity by calculating Bray–Curtis distances via the *phyloseq* package (McMurdie & Holmes, 2013). The Bray–Curtis metric evaluates variations in community composition based on ASV abundances using a permutational multivariate analysis of variance (PERMANOVA) implemented in the *adonis2*() function from the *vegan* package (v 2.6.2; Oksanen et al. 2017). Given that our samples were gathered from five geographical locations, we also examined the influence of locality and incorporated it into our model, setting the “by” parameter to “margin.” As the results for tick species and locality were similar, we chose to present the simpler model that includes locality. Additionally, we utilized *betadispr*(), *betadispr.permutest*(), and post-hoc *TukeyHSD*() to verify that the data satisfies the assumption of homogeneous dispersion for adonis. Finally, we visualized beta diversity through principal coordinate analysis (PCoA), relying on Bray-Curtis distance and using the *phyloseq::ordinate*() function (McMurdie and Holmes, 2014).

### 2.6 Tick microbial networks associations

We investigated microbe–microbe association networks and changes in microbial connectivity between the tick species through quantitative comparisons related to the tick microbiota of *H. lusitanicum* and *R. sanguineus* using NetCoMi (v 1.1.0) (Peschel et al., 2020). Additionally, we analyzed hierarchical clustering of bacterial interactions between *H. lusitanicum* and *R. sanguineus* in localities where they coexist (locations 3 and 4). It’s important to note that *H. lusitanicum* and *R. bursa* were also collected in sympatry at locality 5; however, we could not compare microbial networks there due to the insufficient number of *R. bursa* individuals, which made network comparisons unfeasible. Briefly, we created a single microbial network by agglomerating ASVs into genera using SparCC (Sparse Correlations for Compositional data) (Newman, 2008). This method accounts for the compositional nature of the data through a centered log-ratio transformation (clr) (Zamkovaya et al., 2021; Qannari et al., 2024). To enhance interpretability, we set filter parameters to include only the 50 most frequent taxa and those with over 1000 total reads in the analysis. This reduced the computational nodes in the network analysis. Moreover, we employed the “signed” distance metric, where strong negative correlations increase dissimilarity. We examined network centrality measures, including degree, betweenness, and closeness centrality, which provide insights into the individual roles of taxa within the microbial community. These measures help identify hubs, or *keystone taxa*, as described by Peschel et al. (2020) and that are associated with specific roles in the microbial community. Clusters of nodes, which are closely interconnected yet have few connections outside their module, were identified using greedy modularity optimization provided by the *cluster_fast_greedy* from the *netAnalyze*() function. Hubs were characterized as nodes with an eigenvector centrality value exceeding the empirical 95% quantile of all eigenvector centralities in the network. Additionally, we defined global network properties, including average path length, edge and vertex connectivity, modularity, and clustering coefficient. We visualize the network by assigning colors to determined clusters and scaling node sizes according to the column sums of *clr-transformed* taxa. We investigated changes in the connectivity of microbial association networks between the species *H. lusitanicum* and *R. sanguineus*, while also making quantitative comparisons across the microbial networks of this tick species and also when coexisted in locality 3 and 4, using the *netCompare*() function, implementing a permutation test (setting *permTest* to TRUE) with *nPerm* set to 100 and *nPermRand* to 1000 for the Rand index testing against random cluster assignments for further report the 10 genera exhibiting the highest absolute group differences in degree, betweenness, and closeness centrality.

## 3. Results

### 3.1 Tick microbial community composition

A total of 273 genera and 165 families across 24 phyla were identified in the tick microbiota (**Supplementary file 1**). Proteobacteria was the predominant phylum, accounting for 56% to 4.22% of all ASVs (**Figure 1a-d**). Bacteroidota followed as the second most abundant phylum, with a range of 35% to 12%. This was succeeded by Firmicutes (23% to 7%), Fusobacteriota (23% to 7%), and Actinobacteriota (13% to 0.8%). Additional phyla, such as Synergistota, Spirochaeta, Verrucomicrobiota, Chloroflexi, and Acidobacteriota, are detailed in Supplementary file 1: Dataset S1. At the genus level, the most abundant five genera include *Coxiella* and *Francisella,* which exhibited similar relative abundances ranging from 44% to 0% (**Figure 1a**; **Supplementary file 1: Dataset S3**). Notably, *Coxiella* was more enriched in the tick genera *Rhipicephalus* (*R. sanguineus* and *R. bursa*), while *Francisella* was found to be more prevalent in *H. lusitanicum* ticks. The remaining three genera, *Fusobacterium* (16% to 4.7%), *Porphyromonas* (10% to 3.28%), and *Prevotella* (6.38% to 1.90%) were present in higher proportions specifically within *H. lusitanicum* ticks (**Figure 1c**).

**Figure 1.**
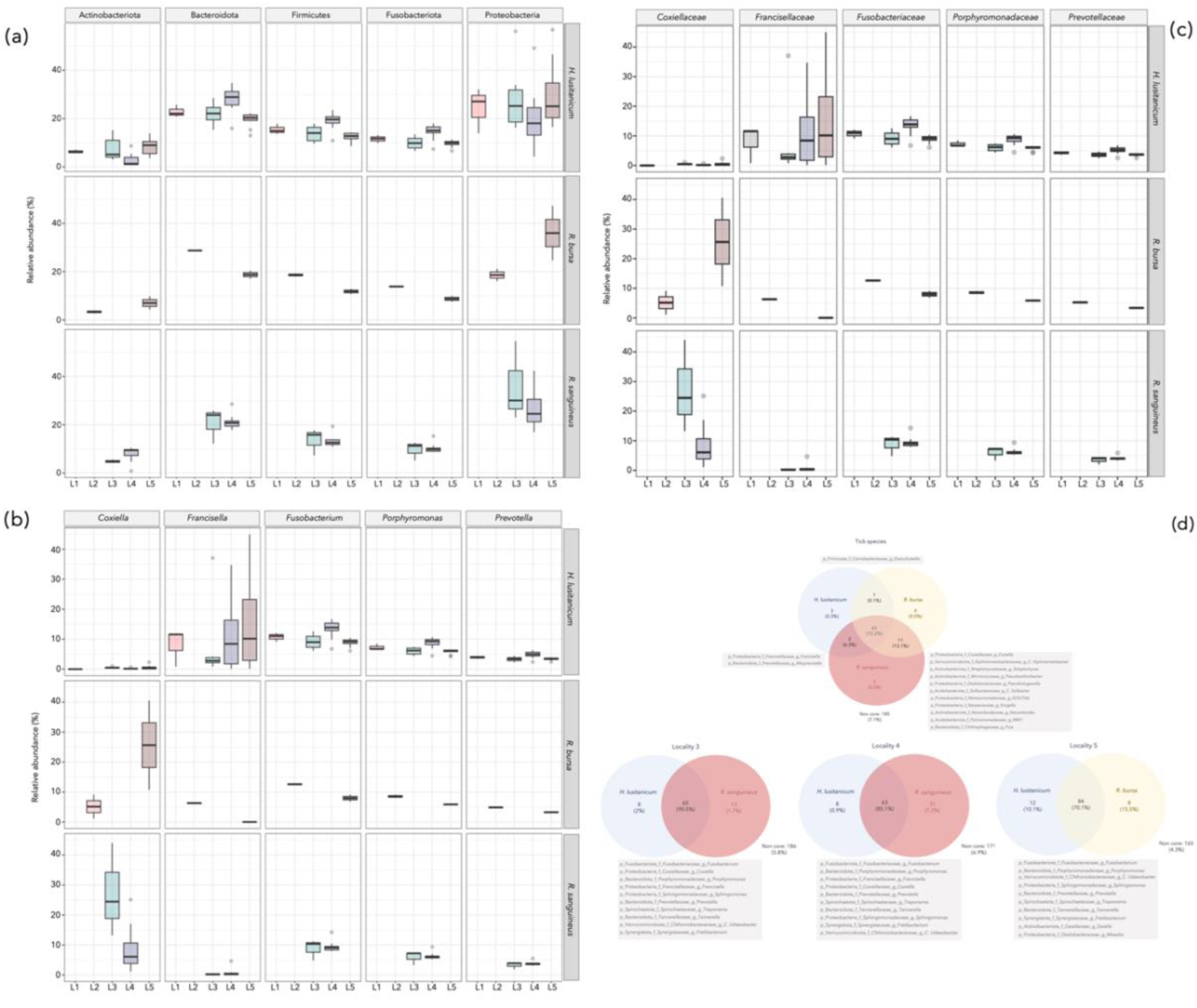
Boxplot showing the top 5 phyla in terms of relative abundance identified within the tick microbiota and grouped by tick species and localities (**Figure 1a**). Boxplot showing the top 10 families and genera in terms of relative abundance identified within the tick microbiota and grouped by tick species and localities (**Figure 1b** and **Figure 1c**, respectively). Venn diagrams depict shared and unique genera among the tick species *H. lusitanicum* and *R. sanguineus*, including those in sympatry at localities 3, 4, and 5. The percentages in brackets represent the average abundance of these genera. Grey boxes indicate shared genera and the ten most abundant shared genera in localities 3, 4, and 5. The complete dataset of core genera identified within and among these tick species is in **Supplementary file 2.**

The Venn diagram illustrated both shared and distinct genera in the tick microbiota based on tick species (host) and their localities (species co-occurring together). In this analysis, we defined the core community as those genera with a relative abundance of at least 0.05% present in at least 85% of the individuals (**Figure 1d**). Among the genera found between tick species, R. bursa and R. sanguineus shared 89% (76 out of 85 genera), while *H. lusitanicum* shared 79% (67 out of 84) and 76% (66 out of 86) with *R. sanguineus* and *R. bursa*, respectively. When analyzing the core microbiota of sympatric tick species, we noted similar findings between *H. lusitanicum* and *R. sanguineus* at locality 3, with 76% (65 out of 86 genera) shared, whereas locality 4 showed a lower sharing rate of 61% (63 out of 102) between these two species. In locality 5, *H. lusitanicum* and *R. bursa* shared 80% of the genera (84 out of 104) in their core microbiota (Figure 1d). Remarkably, the most prevalent shared genera across all three tick species and localities included Fusobacterium (average relative abundance ranging from 13% to 11%) and *Porphyromonas* (also averaging 13% to 11%), followed by *Prevotella, Sphingomonas, Tannerella, Treponema*, and *C. Udaeobacter*, which all had average relative abundances between approximately 5% and 3%. Other notable genera within the core microbiota of the three tick species were *Alistipes, Filifactor, Bacteroides, Streptococcus, Campylobacter, Lactobacillus*, and the *Lachnospiraceae* NK4A136 group, each with an average relative abundance of at least 1% (**Supplementary file 2**). We also noted the *Rickenellaceae* RC9 gut group, which had a lower representation in the core microbiota across the three tick species and locations, averaging around 0.17% (Supplementary file 2). Interestingly, genera like *Coxiella* and *Francisella* were identified as part of the core microbiota only in localities 3 and 4, where *H. lusitanicum* and *R. sanguineus* occurred together, with average relative abundances ranging from 10% to 5% and 7%, respectively (**Supplementary file 2**).

### 3.2 Tick microbiota diversity measures

The alpha diversity indices, namely Shannon, Simpson, and InvSimpson, revealed a significantly diverse microbial community within *H. lusitanicum*, *R. sanguineus*, and *R. bursa* across tick species and genera (**Figures 2 a,b**; **Supplementary File 3: Table S1**). It was observed that no significant differences in Shannon diversity existed, indicating a comparably high richness of the microbial community across the tick species and genera, with low uncertainty in predicting bacterial taxa (Shannon index for tick species; X^2^= 0.43, p-value= 0.8; Shannon index for tick genera; W=234, p-value= 0.7). Similarly, no significant discrepancies were noted in the Simpson indices (Simpson index for tick species; X^2^= 0.22, p-value= 0.89; Simpson index for tick genera; W=258, p-value= 0.8) and InvSimpson indices (InvSimpson index for tick species; W= 0.22, p-value= 0.89; InvSimpson index for tick genera; W=258, p-value= 0.8), further supporting the notion of elevated microbial evenness within both tick species and genera. Consequently, this suggests a high probability of identifying diverse microbial taxa through random sampling of these microbial communities. Hypotheses were examined to elucidate the phylogenetic structure of tick species and genera assemblages. We utilized ses.PD and ses.MPD metrics, which demonstrated no significant differences between the tick species *H. lusitanicum*, *R. sanguineus*, and *R. bursa* (ses.PD, X^2^=0.48, p-value=0.74; ses.MPD, X^2^=0.38, p-value=0.82), nor between the tick genera *Hyalomma* and *Rhipicephalus* (ses.PD, W=226, p-value=0.64; ses.MPD, W=272, p-value=0.56) (**Figures 2c** and **2d; Supplementary File 3: Table S1**). Therefore, it was observed that tick microbial assemblages exhibited a similar pattern at both species and genus levels, appearing clustered in terms of ses.PD (negative ses.PD) while being overdispersed according to ses.MPD (positive ses.MPD) (**Figures 2c-d**).

**Figure 2.**
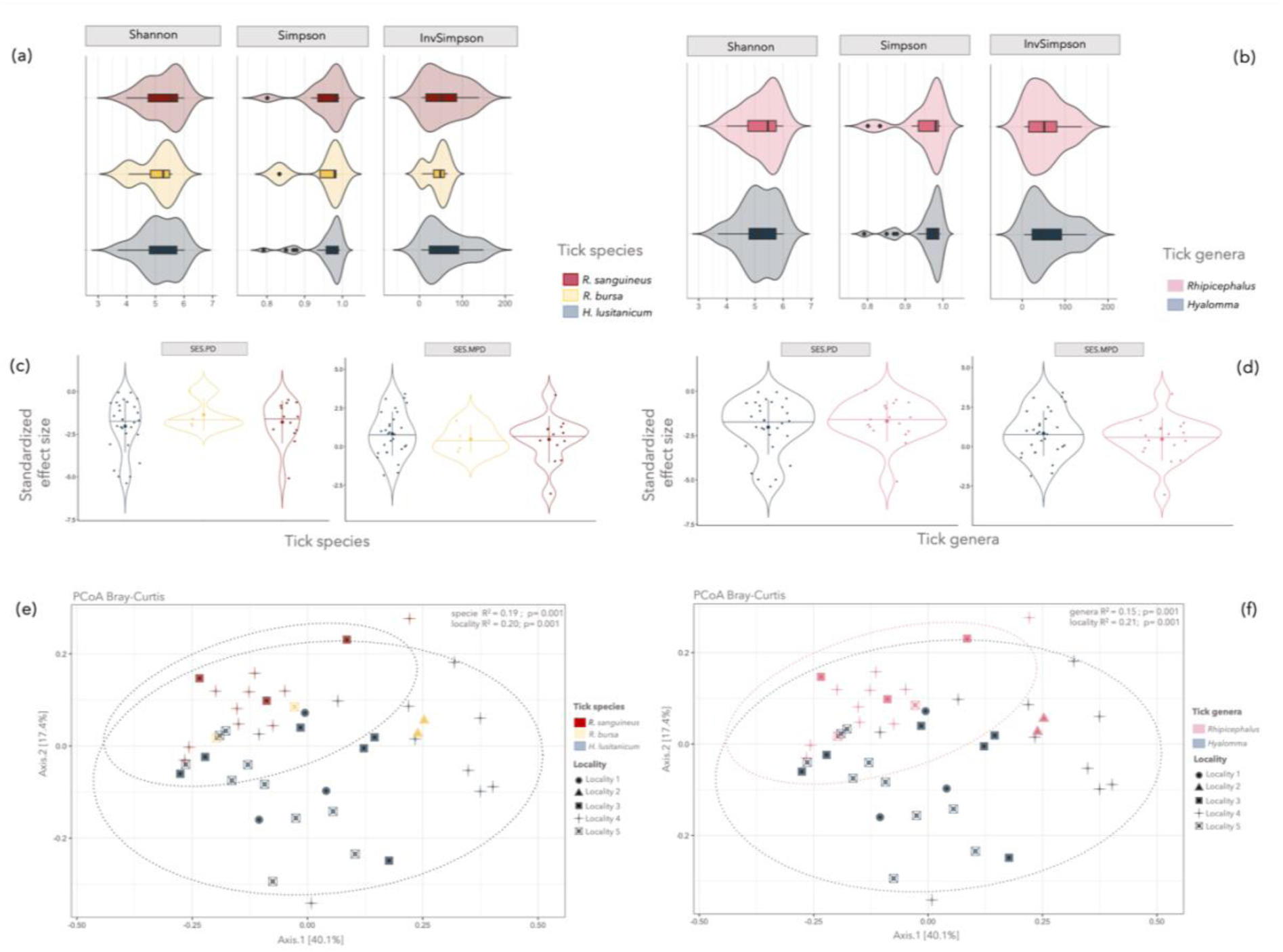
Violin plots illustrate alpha diversity measures (Shannon, Simpson, InvSimpson) across tick species (**Figure 2a**) and genera (**Figure 2b**), using the *violin_plot*() function scaled by width, with horizontal lines denoting quantile density estimates. Phylogenetic metrics SES.PD and SES.MPD are shown for tick species (**Figure 2c**) and genera (**Figure 2d**) via *ggviolin*(), with points shaped by genera, incorporating mean-sd and jittered points. The error plot is a pointrange, with quantiles at 0.5. The complete alpha-diversity estimations are in **Supplementary file 3:Table S1.**

In terms of beta diversity, the analysis indicated that both species and genera might exert a significant influence on the bacterial community composition of tick microbiota. The Bray-Curtis similarity matrix revealed that the structural composition of the tick microbiota significantly varied across species∼locality (PERMANOVA; p=0.001) (Figure 2e) and genera∼locality (PERMANOVA; p=0.001) (**Figure 2f**). The results demonstrated analogous clustering patterns for both species∼locality and genera∼locality, with notable overlap observed between the species *H. lusitanicum* and *R. bursa*, albeit with comparatively greater variation detected between *H. lusitanicum* and *R. sanguineus* (**Figures 2e-f**). As well, the group dispersions of beta diversity calculated with Bray-Curtis distance were not significantly different for the explanatory variable “tick species” (F-value = 1.97, p = 0.1) and “tick genera” (F-value = 0.27, p = 0.6). Thus, any differences detected by the PERMANOVA test can be attributed to differences in their centroid.

### 3.3 Tick microbiota network comparison

To compare networks, the combined count matrix was input into *netConstruct* () along with a binary vector that categorized the samples into one of two groups. The layout derived from the *H. lusitanicum* network was applied to both networks to enhance graphical comparison and highlight distinctions. Additionally, both networks exhibited similar clustering patterns and identified three common hub nodes or keystone taxa: *Fusobacterium, Porphyromonas*, and *Treponema* (**Figure 3a**). A comparison of all global metrics and the ten genera with the largest absolute group differences in degree, betweenness, and closeness centrality can be found in **Supplementary file 3: Tables S2-S3**. Modularity, a property of a network, indicates a proposed division into communities. It assesses the quality of this division based on the number of edges within communities compared to those between them, according to Clauset et al. (20024). Positive modularity values in both networks suggest a potential community structure (Newman, 2006). The clustering coefficient or transitivity of both networks is close to 1 (*H. lusitanicum* = 0.883 and *R. sanguineus* = 0.899), implying that the graphs are nearly fully connected (Kajihara and Hynson 2024). However, no significant differences were observed in the global network metrics (alpha=0.05) (**Supplementary file 3: Table S3**), nor in the centrality measures (**Supplementary file 3: Table S4**). The Jaccard index, which indicates the disparity between the most central node sets in the two groups, is near 1 for closeness, eigenvector centralities, and hub taxa, with a significant probability (P ≥ J), indicating a considerable similarity between the central node sets of both groups (**Supplementary file 3: Table S5**). The Adjusted Rand Index (ARI) shows a significant difference from zero, distinguishing it from random clustering (0.968, p-value=0), thereby indicating a strong resemblance between the two clusterings (**Supplementary file 3: Table S6**).

**Figure 3.**
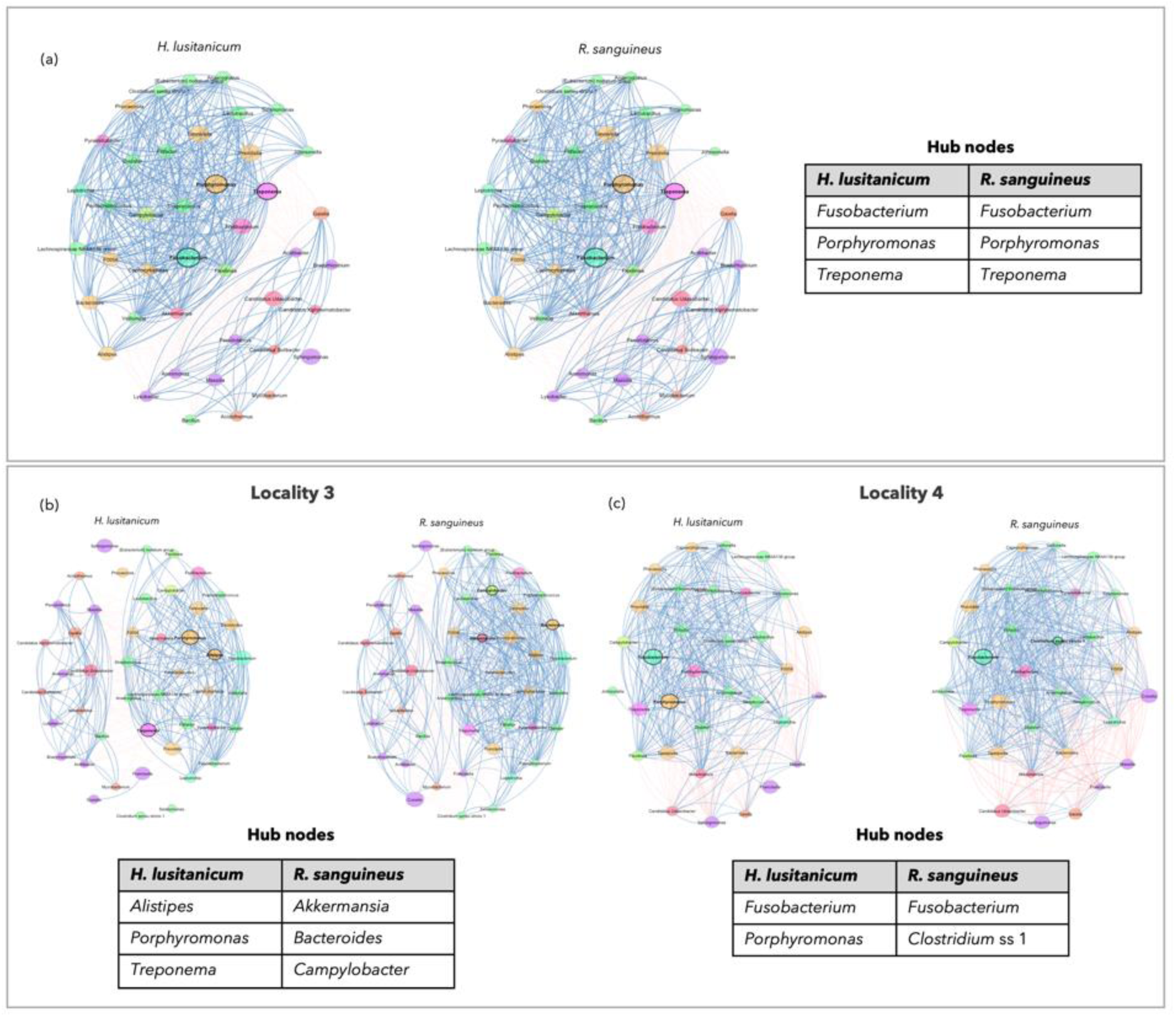
Microbial associations from samples of *H. lusitanicum* and *R. sanguineus* (**Figures 3a**, **3b**, **3c**) were analyzed using the SparCC method in NetCoMi. The signed dissimilarity function converts associations to dissimilarities, with blue edges for positive correlations and red for negative. Node colors indicate phylum, and sizes are clr-transformed. Hubs (in bold) have eigenvector centrality above the 95^th^ percentile. The additional results for whole network properties is in **Supplementary file 3:Table S2-Table S16.**

In locality 3, the sympatric *H. lusitanicum* and *R. sanguineus* networks showed distinct clustering patterns and hub nodes (**Figure 3b; Supplementary file 3: Table S7**). Hub nodes for *H. lusitanicum* include *Alistipes, Porphyromonas*, and *Treponema*, while *Akkermansia, Bacteroides*, and *Campylobacter a*re key taxa in the *R. sanguineus* network (**Supplementary file 3: Tables S8-S9**). In addtion, *H. lusitanicum* has a lower clustering coefficient but higher modularity and positive edge percentage than *R. sanguineus* (**Supplementary file 3: Table S8**). Centrality measure testing showed *R. sanguineus* had a greater degree, with significant differences in closeness centrality between C. *Xiphinematobacter* and *Pseudolabrys* (**Supplementary file 3: Table S9**). The Jaccard index approached 0 for various centralities and hub taxa, indicating distinct central node sets across groups (**Supplementary file 3: Table S10**). While clustering shares some similarities, the Adjusted Rand Index (ARI) differs significantly from zero, with an ARI of 0.659 (p-value= 0), indicating a high level of similarity between clusterings (**Supplementary file 3: Table S11**).

In comparing the networks of sympatric *H. lusitanicum* and *R. sanguineus* at locality 4, a different clustering pattern emerged, similar to locality 3, except for one shared hub node: *Fusobacterium* (**Figure 3c; Supplementary file 3: Table S12**). Detailed global measures and the ten genera with the highest absolute group differences in degree, betweenness, and closeness centrality are in **Supplementary file 3: Tables S13-S14**. Significant differences appeared only in positive edge percentages for centrality measures, with *R. sanguineus* at 73.81% and *H. lusitanicum* at 65.92%. None of the measures for the ten genera showed significance (**Supplementary file 3: Table S14**). The Jaccard index nearing 1 suggests significant probability (P≥ J) for eigenvector centralities; however, low hub node values indicate only one shared node (*Fusobacterium*) (**Supplementary file 3: Table S15**). The Adjusted Rand Index (ARI) significantly deviates from zero, indicating non-random clustering (0.220, p-value = 0) (Supplementary file 3: Table S16). A consistent pattern in all three analyses showed that *H. lusitanicum* tends to have a lower clustering coefficient, higher modularity, and greater positive edge percentage than *R. sanguineus* (even if not significant) across tested hypotheses, except at locality 4, where *R. sanguineus* had a higher positive edge percentage (**Supplementary file 3: Tables S3; S8; S13**).

## 4. Discussion

This study examined the microbial communities associated with three wild tick species—*H. lusitanicum, R. sanguineus*, and *R. bursa*—collected in various locations throughout Castilla-La Mancha, Spain. The sympatric presence of these species in certain localities demonstrated ecological interactions and shared environmental factors. Previous research has often documented these tick species in the same regions, feeding on Eurasian wild boar (*Sus scrofa*) and red deer (*Cervus elaphus*) (Fernández de Mera et al., 2013). Notably, *H. lusitanicum* is extensively distributed in central Spain, making up over 80% of the observed tick community (Requena-García et al., 2017; Valcárcel et al., 2014). We then investigated the microbiota composition of these three species, focusing on both shared characteristics and unique traits that may affect vector competence and ecological dynamics roles. At the phylum level, Proteobacteria were predominant in all three tick species, consistent with earlier studies conducted in Spain and worldwide (Narasimhan and Fikrig, 2015; Portillo et al., 2019; Díaz-Sánchez et al., 2021). Other phyla, such as Bacteroidota, Firmicutes, Actinobacteriota, and Fusobacteriota, were also significant, highlighting the diversity of tick-associated microbiota. Noteworthy findings at the genus level include the predominance of *Coxiella* and *Francisella*. These genera exhibit a range of behaviors that encompass critical endosymbiotic functions essential for the survival of ticks, as well as pathogens implicated in diseases such as Q fever (*Coxiella burnetii*) and tularemia (*Francisella tularensis*) (Gerhart et al., 2016; Andreotti et al., 2011; Ben-Yosef et al., 2020; Duron et al., 2015; Duron et al., 2017). The presence of the genus *Francisella* in *H. lusitanicum* and *Coxiella* in *R. sanguineus* and *R. bursa* is consistent with earlier findings (Toledo et al., 2009; Díaz-Sánchez et al., 2021; Ben-Yosef et al., 2020; Herrera et al., 2023), suggesting that evolutionary factors may drive these microbial associations. Additionally, we observed a significant community of shared genera among tick species, pointing to a likely well-conserved core bacteria **(Figure 1d**). Xu et al. (2024) previously revealed that shared taxa represent a major part of tick bacterial microbiota. Prior research has shown that ticks in close geographical proximity tend to share microbes (Van Treuren et al., 2015; Portillo et al., 2019). This pattern can also be facilitated by co-feeding behaviors, where two or more ticks share microbes by feeding closely together on the same host (Aivelo et al., 2022). Among the shared genera, *Fusobacterium, Porphyromonas*, and *Prevotella* were the most prevalent. *Fusobacterium* is a widespread bacterium found in soil, manure, the human gastrointestinal tract, as well as on the skin and hooves of domestic animals. It has also been identified as a commensal in tick microbiota (Delano et al., 2002; Fountain-Jones et al., 2023). *Porphyromonas* has been detected in soil and within the intestinal microbiota of arthropods (Acuña-Amador and Barloy-Hubler, 2020). Numerous microorganisms discovered are linked to the soil in ecosystems where these arthropods thrive, emphasizing the role of environmental microbes in shaping tick microbial communities. (Portillo et al., 2019; Narasimhan et al., 2021). *Prevotella* was observed in *H. lusitanicum, I. persulcatus* (Zhang et al., 2014; Díaz-Sánchez et al., 2021), as well as in the hemiptera, Triatominae (commonly known as kissing bugs) (Montoya-Porras et al., 2018). The potential functional roles of these genera, including *Prevotella* contribution to blood digestion and *Fusobacterium* role in environmental nutrient cycling, merit further exploration to clarify their impact on tick biology and their competence as secondary symbionts (Folk and Leung, 2002). A general understanding of tick-bacteria microbiota and how OTU/ASV abundance affects functionality is still limited. While abundant OTUs are often viewed as functionally significant, it is crucial to recognize that rare OTUs/ASVs can also have a substantial effect on interactions between the host and microbes (Lozupone et al., 2012; Aivelo et al., 2022).

Similar microbial diversity (alpha-diversity) was noted among *H. lusitanicum, R. sanguineus*, and *R. bursa* (refer to Figure 2a, b). Analysis of phylogenetic metrics revealed clustering patterns and overdispersion, suggesting that deterministic processes such as competitive exclusion and habitat filtering influence the microbial communities associated with these three tick species. The observed metrics mismatches appear to stem from an overdispersion of clusters, which relates to basal overdispersion and terminal clustering (Goberna et al., 2014a,b; Mazel et al., 2016). These results align with the patterns of “basal” overdispersion, where the co-occurrence of distantly related bacteria reflects host-specific processes structuring the bacterial assemblages, potentially favoring certain bacterial lineages (Bevins & Salzman, 2011; Hooper et al., 2012; Mazel et al., 2016). Consequently, microbial competition-driven coexistence may produce phylogenetic overdispersion patterns, suggesting that distantly related species with minimal niche overlap and competition are more likely to coexist. In contrast, phylogenetic clustering is believed to signify the coexistence of closely related species sharing a similar environmental niche (Mazel et al., 2016). A prior study by Díaz-Sánchez et al. (2019) found overdispersion patterns using MDP metrics in *Ixodes affinis, I. scapularis, I. ricinus*, and *I. ovatus* microbial communities, while *I. persulcatus, I. pavloskyi,* and *I. ventalloi* exhibited phylogenetic clustering. Although these findings are intriguing, further studies should explore the connection between these patterns and the ecological processes and whether they may relate to a continuum of abiotic and biotic factors affecting selection in the tick microbial community assembly. At present, we can only speculate regarding the processes potentially involved in tick microbial assembly. For instance, the haematophagous behavior of ticks may necessitate a complex digestive mechanism that requires various biochemical compounds for the degradation process (Mans and Neitz, 2004). This may lead to the emergence of distinct bacterial clades that possess unique enzymatic profiles within the tick microbiota, likely resulting in overdispersion patterns characterized by the co-occurrence of distantly related lineages. Within these clades, host-specific processes like mucus barriers, oxygen levels, and the tick immune system may favor certain lineages across a broad phylogenetic scale. Indeed, feeding from a single host individual at each life stage can enhance opportunities for deterministic processes in microbial assembly (Aivelo et al., 2022). The analysis of beta-diversity metrics demonstrated that both host-specific and ecological factors significantly influence tick microbiota composition. This finding highlights the importance of conducting longitudinal studies to track microbial dynamics across various tick species and environmental gradients. Notably, the visualization of beta-diversity illustrates the presence of overlapping microbial communities among different tick species, particularly between *H. lusitanicum* and *R. bursa*. This overlap may suggest shared environmental conditions, as ticks residing in similar geographic areas might harbor a similar microbiota (Portillo et al., 2019; Xu et al., 2023). A similar pattern was observed by Herrera et al. (2023), who reported a corresponding pattern of beta-diversity in ticks from Castilla-Leon (northwestern Spain), where *R. bursa* and *R. sanguineus* clustered similarly with *H. marginatum*.

Microbial co-occurrence networks identified key taxa such as *Fusobacterium, Porphyromonas*, and *Treponema*, consistently found across different tick species and locations in this study. The significance of *Fusobacterium* as a central taxon has been previously noted as an ecological indicator for pathogens like *Anaplasma* (Fountain-Jones et al., 2023). Variations in network characteristics, including clustering coefficients and modularity, indicate species-specific and locality-specific factors that may affect microbial network relationships. Further research is needed to gain more understanding of the biological implications of these findings and how pathogens and symbionts interact with tick species, with environmental microbes influencing these dynamics. A study by Xu et al. (2023) revealed that environmental microorganisms are pivotal in microbial co-occurrence networks, displaying fluctuating patterns among various groups. This aligns with our findings, suggesting that soil microbes could impact the composition of tick microbiota (Naramsiham and Fikrig, 2015; Kwan et al., 2017). Literature indicates that *Porphyromonas* has been detected in the intestinal microbiota of arthropods, including ectoparasites and insects, and has been linked to human-altered soils (agricultural land and crops) (Acuña-Amador and Hubler, 2020). *Treponema* species, which are obligate parasites found in a wide range of animal hosts, have some basal species that might be free-living (Norris et al., 2006). *Treponema* can be both pathogenic and non-pathogenic, sometimes playing a symbiotic role in hosts; for instance, in termite guts, they contribute to H_2_-CO_2_ acetogenesis and nitrogen fixation, supplying carbon and energy in return for nitrogen (Graber et al., 2004; Gogarten et al., 2016). In *H. lusitanicum*, we noted a slightly lower clustering coefficient, alongside higher modularity and positive edge percentage compared to *R. sanguineus*, although these differences were not statistically significant across all tested hypotheses. In tick microbial networks, positive associations are significantly more prevalent than negative ones (Aivelo et al., 2019; Lejal et al., 2021; Fountain-Jones et al., 2023). Fountain-Jones et al. (2023) emphasized the vital importance of beneficial microbial relationships in influencing the microbiota of *I. scapularis*. The researchers discovered that positive interactions are common, implying a facilitation process and a trend of simultaneous colonization. Consequently, it is plausible that ticks may acquire multiple microbial groups at once. A positive edge percentage may suggest enhanced connectivity among nodes, which could increase the risk of transmitting an external stressor throughout the ecosystem, potentially harming the host (Kahijara and Hynson, 2024). These network analyses elucidate the intricate relationships between environmental microbes and tick microbiota, shedding light on possible pathways for pathogen acquisition and transmission. Quantifying associations in natural settings and their relative significance against other ecological drivers presents a methodological challenge. Accounting for spatial and temporal patterns is essential, as species may co-occur less often than expected by chance due to spatial segregation or limitations in dispersal rather than microbial competition (Fountain-Jones et al., 2023). Sympatric tick species may show differences in pathogen competence stemming from variations in host preferences, physiology, and pathogen specificity. Additionally, co-infection dynamics within these populations can influence vector competence through interactions among pathogens.

## 5. Conclusions

This study demonstrates a notable richness and diversity of bacterial microbiota associated with wild ticks, indicative of a stable core bacteria, although interesting divergences among tick species may highlight the specificity of their microbiota. The findings pave the way for further research on how functional microbiota shape tick biology, particularly regarding shared and species-specific microbial assemblages. Future efforts should focus on 1) longitudinal studies to assess the temporal stability of core tick microbiota; 2) experimental research on essential genera like *Francisella* and *Coxiella*, as well as on complex interactions between environmental microbes and tick microbiota, highlighting potential pathways for pathogen acquisition and transmission; 3) integrative methods that combine ecological, genetic, and environmental data to reveal factors influencing microbial community assembly; and 4) should focus on advancing the understanding of tick-pathogen and tick-host interactions on microbiota composition. Ultimately, this work underscores the importance of microbiota in tick vector capacity, opening avenues for microbiota-focused interventions in vector control.

## CRediT Author Statement

**Victor Noguerales:** Conceptualization, investigation, formal analysis, methodology writing-original draft, review and editing. **José de la Fuente:** Review and editing. **Sandra Díaz-Sánchez:** Conceptualization, investigation, formal analysis, methodology writing-original draft, review, editing and funding acquisition.

