## Supplementary material for "Understanding the mechanisms underlying microbiota variation in wild tick populations": Supplementary file 3.pdf

**Table S1.** Alpha diversity estimations within tick samples shown in **Figure 2a, b, c,** and **d**.

| | | | | non-phylogenetic diversity index | | | phylogenetic diversity index | |
|---|---|---|---|---|---|---|---|---|
| **ID samples** | **Tick genera** | **Tick species** | **Locality** | **Shannon** | **Simpson** | **InvSimpson** | **ses.PD** | **ses. MPD** |
| T1L20 | Hyalomma | *H. lusitanicum* | Locality 1 | 5.07 | 0.97 | 37.37 | -0.05 | 1.07 |
| T1L21 | Hyalomma | *H. lusitanicum* | Locality 1 | 5.11 | 0.97 | 29.66 | -1.82 | 1.23 |
| T1L22 | Hyalomma | *H. lusitanicum* | Locality 1 | 5.76 | 0.99 | 95.77 | -2.14 | 0.93 |
| T1L64 | Rhipicephalus | *R. bursa* | Locality 2 | 5.46 | 0.98 | 66.30 | -1.59 | 0.69 |
| T1L65 | Rhipicephalus | *R. bursa* | Locality 2 | 5.09 | 0.98 | 42.49 | -2.01 | -0.10 |
| T1L55 | Rhipicephalus | *R. sanguineus* | Locality 3 | 5.17 | 0.97 | 38.72 | -0.97 | 1.17 |
| T1L56 | Rhipicephalus | *R. sanguineus* | Locality 3 | 3.99 | 0.80 | 5.02 | -1.83 | 0.10 |
| T1L57 | Rhipicephalus | *R. sanguineus* | Locality 3 | 4.75 | 0.93 | 14.95 | -2.22 | 0.39 |
| T1L58 | Hyalomma | *H. lusitanicum* | Locality 3 | 6.02 | 0.99 | 148.82 | -0.82 | -0.38 |
| T1L59 | Hyalomma | *H. lusitanicum* | Locality 3 | 5.94 | 0.99 | 149.03 | -0.64 | -1.86 |
| T1L60 | Hyalomma | *H. lusitanicum* | Locality 3 | 5.18 | 0.98 | 40.13 | -1.43 | 1.00 |
| T1L61 | Hyalomma | *H. lusitanicum* | Locality 3 | 3.82 | 0.85 | 6.69 | -1.62 | 1.45 |
| T1L62 | Hyalomma | *H. lusitanicum* | Locality 3 | 4.89 | 0.95 | 20.80 | -1.65 | 1.33 |
| T1L63 | Hyalomma | *H. lusitanicum* | Locality 3 | 5.58 | 0.99 | 91.68 | -2.08 | 0.84 |
| T1L27 | Rhipicephalus | *R. sanguineus* | Locality 4 | 4.65 | 0.92 | 12.03 | -2.91 | 1.29 |
| T1L28 | Rhipicephalus | *R. sanguineus* | Locality 4 | 5.75 | 0.99 | 86.85 | -1.47 | -3.05 |
| T1L29 | Rhipicephalus | *R. sanguineus* | Locality 4 | 5.55 | 0.98 | 63.15 | -0.82 | 1.12 |
| T1L30 | Rhipicephalus | *R. sanguineus* | Locality 4 | 4.96 | 0.93 | 15.33 | -0.63 | 1.50 |
| T1L39 | Rhipicephalus | *R. sanguineus* | Locality 4 | 5.81 | 0.99 | 119.70 | -2.26 | 0.89 |
| T1L40 | Rhipicephalus | *R. sanguineus* | Locality 4 | 5.78 | 0.99 | 80.66 | -2.44 | -0.89 |
| T1L41 | Rhipicephalus | *R. sanguineus* | Locality 4 | 6.01 | 0.99 | 139.14 | -0.49 | -0.95 |
| T1L42 | Rhipicephalus | *R. sanguineus* | Locality 4 | 5.59 | 0.98 | 51.56 | -0.67 | 0.80 |
| T1L43 | Rhipicephalus | *R. sanguineus* | Locality 4 | 5.83 | 0.99 | 114.64 | -1.47 | 0.36 |
| T1L45 | Rhipicephalus | *R. sanguineus* | Locality 4 | 4.55 | 0.96 | 22.92 | -5.08 | 3.34 |
| T1L44 | Hyalomma | *H. lusitanicum* | Locality 4 | 4.58 | 0.96 | 23.22 | -4.98 | 2.25 |
| T1L46 | Hyalomma | *H. lusitanicum* | Locality 4 | 4.60 | 0.96 | 26.15 | -5.37 | 1.76 |
| T1L47 | Hyalomma | *H. lusitanicum* | Locality 4 | 4.79 | 0.97 | 35.51 | -3.14 | 2.51 |
| T1L48 | Hyalomma | *H. lusitanicum* | Locality 4 | 4.56 | 0.95 | 20.30 | -4.98 | 3.09 |
| T1L49 | Hyalomma | *H. lusitanicum* | Locality 4 | 4.95 | 0.98 | 47.13 | -4.63 | 2.24 |
| T1L50 | Hyalomma | *H. lusitanicum* | Locality 4 | 5.06 | 0.98 | 52.96 | -4.20 | 2.84 |
| T1L51 | Hyalomma | *H. lusitanicum* | Locality 4 | 5.17 | 0.98 | 57.18 | -2.52 | 3.41 |
| T1L52 | Hyalomma | *H. lusitanicum* | Locality 4 | 5.77 | 0.99 | 109.20 | -2.08 | -1.18 |
| T1L53 | Hyalomma | *H. lusitanicum* | Locality 4 | 4.14 | 0.87 | 7.70 | -0.90 | -0.16 |
| T1L54 | Hyalomma | *H. lusitanicum* | Locality 4 | 5.51 | 0.99 | 78.23 | -3.09 | -0.70 |
| T1L23 | Hyalomma | *H. lusitanicum* | Locality 5 | 5.03 | 0.96 | 23.57 | -0.41 | 3.16 |
| T1L24 | Hyalomma | *H. lusitanicum* | Locality 5 | 4.89 | 0.93 | 14.76 | -1.68 | 0.08 |
| T1L26 | Hyalomma | *H. lusitanicum* | Locality 5 | 3.68 | 0.79 | 4.79 | -2.45 | 0.49 |
| T1L31 | Hyalomma | *H. lusitanicum* | Locality 5 | 4.41 | 0.88 | 8.20 | -0.46 | 0.37 |
| T1L32 | Hyalomma | *H. lusitanicum* | Locality 5 | 6.01 | 0.99 | 149.36 | -0.06 | -1.69 |
| T1L33 | Hyalomma | *H. lusitanicum* | Locality 5 | 5.50 | 0.98 | 45.12 | -0.61 | 0.79 |
| T1L35 | Hyalomma | *H. lusitanicum* | Locality 5 | 5.71 | 0.99 | 79.62 | -1.24 | 0.00 |
| T1L36 | Hyalomma | *H. lusitanicum* | Locality 5 | 5.93 | 0.99 | 125.71 | -2.02 | 0.12 |
| T1L37 | Hyalomma | *H. lusitanicum* | Locality 5 | 5.75 | 0.99 | 76.59 | -0.67 | -0.42 |
| T1L38 | Hyalomma | *H. lusitanicum* | Locality 5 | 5.96 | 0.99 | 134.41 | -1.11 | -0.36 |
| T1L34 | Rhipicephalus | *R. bursa* | Locality 5 | 5.59 | 0.98 | 55.28 | -1.89 | -0.32 |
| T1L25 | Rhipicephalus | *R. bursa* | Locality 5 | 4.06 | 0.83 | 5.99 | 0.05 | 1.67 |

### Additional results for whole network properties of *H. lusitanicum* and *R. sanguineus*

**Table S2.** Properties of the networks shown in **Figure 3a**: a) Frequency table of clusters in the tick species *H. lusitanicum* network. b) Frequency table of clusters in the tick species *R. sanguineus* network. c) Detected hub nodes in both groups. d)-f) Centrality values of the genera with the highest centrality in decreasing order. The upper part of the table contains the five genera with the highest centrality in *H. lusitanicum* and the lower part with the highest centrality in *R. sanguineus* respectively. Thus, a genus can occur twice in the same table (Peschet et al., 2020).

a) Cluster at *H. lusitanicum*

| Name | Frequency |
|---|---|
| 0 | 3 |
| 1 | 28 |
| 2 | 13 |

b) Cluster at *R. sanguineus*

| Name | Frequency |
|---|---|
| 0 | 2 |
| 1 | 28 |
| 2 | 14 |

c) Hub nodes, in alphabetical order. Based on empirical quantiles of centralities.

| *H. lusitanicum* | *R. sanguineus* |
|---|---|
| *Fusobacterium* | *Fusobacterium* |
| *Porphyromonas* | *Porphyromonas* |
| *Treponema* | *Treponema* |

d) Degree

| Genus | | |
|---|---|---|
| Highest values on the *H. lusitanicum* group: | *H. lusitanicum* | *R. sanguineus* |
| *Fretibacterium* | 0.81395 | 0.90698 |
| *Campylobacter* | 0.81395 | 0.90698 |
| *Filifactor* | 0.81395 | 0.86047 |
| *Dialister* | 0.81395 | 0.88372 |
| *Streptococcus* | 0.7907 | 0.88372 |
| Highest values on the *R. sanguineus* group: | *H. lusitanicum* | *R. sanguineus* |
| *Fretibacterium* | 0.81395 | 0.90698 |
| *C. Udaeobacter* | 0.72093 | 0.90698 |
| *Campylobacter* | 0.81395 | 0.90698 |
| *Gaiella* | 0.7907 | 0.90698 |
| *Massilia* | 0.72093 | 0.90698 |

e) Betweennes centrality

| Genus | | |
|---|---|---|
| Highest values on the *H. lusitanicum* group: | *H. lusitanicum* | *R. sanguineus* |
| *Fretibacterium* | 0.06538 | 0.04024 |
| *Gaiella* | 0.05385 | 0.00854 |
| *Arenimonas* | 0.03846 | 0.00976 |
| *Massilia* | 0.03205 | 0.02073 |
| *Acidibacter* | 0.02692 | 0 |
| Highest values on the *R. sanguineus* group: | *H. lusitanicum* | *R. sanguineus* |
| *Fretibacterium* | 0.06538 | 0.04024 |
| C. *Solibacter* | 0 | 0.03293 |
| *Streptococcus* | 0 | 0.02683 |
| *Massilia* | 0.03205 | 0.02073 |
| C. *Udaeobacter* | 0.01538 | 0.01341 |

f) Eigenvector centrality

| Genus | | |
|---|---|---|
| Highest values on the *H. lusitanicum* group: | *H. lusitanicum* | *R. sanguineus* |
| Fusobacterium | 1 | 1 |
| Porphyromonas | 0.99074 | 0.99871 |
| Treponema | 0.9905 | 0.99223 |
| Filifactor | 0.98895 | 0.96289 |
| Tannerella | 0.98618 | 0.99111 |
| Highest values on the *R. sanguineus* group: | *H. lusitanicum* | *R. sanguineus* |
| Fusobacterium | 1 | 1 |
| Porphyromonas | 0.99074 | 0.99871 |
| Treponema | 0.9905 | 0.99223 |
| Tannerella | 0.98618 | 0.99111 |
| Streptococcus | 0.94804 | 0.99071 |

**Table S3.** Comparison of network properties. Results from testing global network metrics of the networks shown in **Figure 3a** for group differences (via a permutation procedure using 1000 permutations). The computed measures for *H. lusitanicum* and *R. sanguineus*, the absolute difference, and the p-value for testing the null hypothesis are shown, respectively, as H 0: |diff| = 0. Global network properties are defined for the whole network an offer an insight into the overall network structure (Peschet el at., 2020).

| **Whole network** | | | | |
|---|---|---|---|---|
| | *H. lusitanicum* | *R. sanguineus* | abs. difference | p-value |
| Number of components | 4.000 | 3.000 | 1.000 | 0.712871 |
| Clustering coefficient | 0.883 | 0.899 | 0.016 | 0.811881 |
| Modularity | 0.054 | 0.027 | 0.027 | 0.396040 |
| Positive edge percentage | 68.850 | 58.126 | 10.724 | 0.069307 |
| Edge density | 0.597 | 0.722 | 0.125 | 0.366337 |
| Natural connectivity | 0.424 | 0.402 | 0.021 | 0.792079 |

p-values: one-tailed test with null hypothesis diff=0

**Table S4.** Results from testing centrality measures of the networks in **Figure 3a** for group differences (via a permutation procedure using 1000 permutations). Shown are, respectively, the computed measures for *H. lusitanicum* and *R. sanguineus*, the absolute difference, and the p-value for testing the null hypothesis H 0: |diff| = 0 (Peschet et al., 2020). The p-values are adjusted for multiple testing using the adaptive Benjamini-Hochberg method (Benjamini and Hochberg, 2000), where the proportion of true H 0 is determined according to Langaas et al. (2005). These results contain 10 subjects with the highest absolute group difference. All measures are normalized to [0,1].

| | *H. lusitanicum* | *R. sanguineus* | abs. difference | p-value |
|---|---|---|---|---|
| **Degree** (normalized) | | | | |
| *Mycobacterium* | 0.047 | 0.791 | 0.744 | 0.625611 |
| *Bacillus* | 0.279 | 0.907 | 0.628 | 0.607211 |
| *Bradyrhizobium* | 0.093 | 0.698 | 0.605 | 0.885516 |
| *Sphingomonas* | 0.000 | 0.558 | 0.558 | 0.485769 |
| *Lachnospiraceae NK4A136 group* | 0.767 | 0.279 | 0.488 | 0.625611 |
| *Candidatus Solibacter* | 0.209 | 0.698 | 0.488 | 0.520467 |
| *Johnsonella* | 0.349 | 0.047 | 0.302 | 0.885516 |
| *Fusobacterium* | 0.628 | 0.884 | 0.256 | 0.485769 |
| *Porphyromonas* | 0.628 | 0.884 | 0.256 | 0.485769 |
| *Treponema* | 0.628 | 0.884 | 0.256 | 0.485769 |
| **Betweenness centrality** (normalized) | | | | |
| *Gaiella* | 0.054 | 0.009 | 0.045 | 0.925743 |
| Candidatus *Solibacter* | 0.000 | 0.033 | 0.033 | 0.925743 |
| *Arenimonas* | 0.038 | 0.010 | 0.029 | 0.925743 |
| *Acidibacter* | 0.027 | 0.000 | 0.027 | 0.925743 |
| *Streptococcus* | 0.000 | 0.027 | 0.027 | 0.925743 |
| *Fretibacterium* | 0.065 | 0.040 | 0.025 | 0.925743 |
| *Fusobacterium* | 0.022 | 0.005 | 0.017 | 1.000000 |
| *Dialister* | 0.017 | 0.000 | 0.017 | 0.925743 |
| *Filifactor* | 0.013 | 0.000 | 0.013 | 0.925743 |
| *Massilia* | 0.032 | 0.021 | 0.011 | 0.925743 |
| **Closeness centrality** (normalized) | | | | |
| Candidatus *Xiphinematbacter* | 0.742 | 2.029 | 1.287 | 0.292661 |
| Candidatus *Solibacter* | 0.928 | 2.167 | 1.238 | 0.292661 |
| *Sphingomonas* | 0.000 | 1.086 | 1.086 | 0.292661 |
| *Pseudolabrys* | 1.107 | 2.085 | 0.978 | 0.292661 |
| *Acidothermus* | 1.087 | 1.941 | 0.854 | 0.292661 |
| *Gaiella* | 1.261 | 2.042 | 0.782 | 0.292661 |
| *Bacillus* | 0.894 | 1.674 | 0.780 | 0.292661 |
| *Massilia* | 1.324 | 2.102 | 0.778 | 0.292661 |
| *Mycobacterium* | 0.736 | 1.395 | 0.659 | 0.600329 |
| *Bradyrhizobium* | 0.795 | 1.403 | 0.608 | 0.600329 |
| Significance codes: ∗∗∗ : 0.001, ∗∗: 0.01, ∗:0.05, .: 0.1 | | | | |

**Table S5**. Jaccard index values express the similarity of the sets of most central nodes and the sets of hub taxa between the two networks to assess how different the two sets of most central nodes are between the groups. Jaccard's index is 0 if the sets are completely different and 1 for exactly equal sets. P(J ≤ j) is the probability that Jaccard's index takes for the present total number of taxa a value less than or equal to the calculated index j (P(J ≥ j) is defined analogously) (Peschel et al., 2020).

| | Jacc | P(J ≤ Jacc) | P(J ≥ Jacc) |
|---|---|---|---|
| **degree** | 0.250 | 0.468221 | 0.804908 |
| **betweenness centr.** | 0.33 | 0.618372 | 0.595935 |
| **closeness centr.** | 0.833 | 0.999953 | 0.000544 *** |
| **eigenvec. centr.** | 0.833 | 0.999953 | 0.000544 *** |
| **hub taxa** | 1.000 | 1.000000 | 0.037037 * |
| Significance codes:∗∗∗ : 0.001, ∗∗: 0.01, ∗:0.05, .: 0.1 | | | |

**Table S6.** Adjusted Rand index assessed similarity between clusterings and range between -1 and 1, where 1 corresponds to identical clustering and 0 to the expected value for two random clusterings. Consequently, positive values imply that two clusters are more similar, and negative values are less similar than expected at random (Peschel et al., 2020).

| Adjusted Rand index | |
|---|---|
| | **Whole network** |
| **ARI** | 0.968 |
| **p-value** | 0.000 |

ARI in [-1,1] with ARI=1: perfect agreement between clusterings; ARI=0: expected for two random clusterings
p-value: permutation test (n=1000) with null hypothesis ARI=0

### Additional results for whole network properties of *H. lusitanicum* and *R. sanguineus* collected at locality 3

**Table S7**. Properties of the networks shown in **Figure 3b**: a) Frequency table of clusters in the tick species *H. lusitanicum* network in locality 3. b) Frequency table of clusters in the tick species R. sanguineus network in locality 3. c) Detected hub nodes in both groups. d)-f) Centrality values of the genera with the highest centrality in decreasing order. The upper part of the table contains the five genera with the highest centrality in *H. lusitanicum* and the lower part with the highest centrality in *R. sanguineus* respectively. Thus, a genus can occur twice in the same table (Peschet el at., 2020).

a) Cluster at *H. lusitanicum*

| Name | Frequency |
|---|---|
| 0 | 4 |
| 1 | 15 |
| 2 | 25 |
| 3 | 2 |

b) Cluster at *R. sanguineus*

| Name | Frequency |
|---|---|
| 1 | 30 |
| 2 | 16 |

c) Hub nodes, in alphabetical order. Based on empirical quantiles of centralities.

| *H. lusitanicum* | *R. sanguineus* |
|---|---|
| *Alistipes* | *Akkermansia* |
| *Porphyromonas* | *Bacteroides* |
| *Treponema* | *Campylobacter* |

d) Degree

| Genus | | |
|---|---|---|
| Highest values on the *H. lusitanicum* group: | *H. lusitanicum* | *R. sanguineus* |
| Porphyromonas | 0.6667 | 0.75556 |
| Faecalibacterium | 0.64444 | 0.82222 |
| Alistipes | 0.62222 | 0.68889 |
| Treponema | 0.57778 | 0.71111 |
| Candidatus Udeobacter | 0.57778 | 0.84444 |
| Highest values on the *R. sanguineus* group: | *H. lusitanicum* | *R. sanguineus* |
| Massilia | 0.55556 | 0.86667 |
| Candidatus Udeobacter | 0.57778 | 0.84444 |
| Dialister | 0.42222 | 0.84444 |
| Capnocytophaga | 0 | 0.84444 |
| Arenimonas | 0.42222 | 0.84444 |

e) Betweenness centrality

| Genus | | |
|---|---|---|
| Highest values on the *H. lusitanicum* group: | *H. lusitanicum* | *R. sanguineus* |
| Porphyromonas | 0.05803 | 0.10303 |
| C. Xiphinematobacter | 0.05668 | 0.0303 |
| Alistipes | 0.05398 | 0.08485 |
| Faecalibacterium | 0.05398 | 0.1202 |
| Treponema | 0.04588 | 0.01616 |
| Highest values on the *R. sanguineus* group: | *H. lusitanicum* | *R. sanguineus* |
| Dialister | 0.00405 | 0.15253 |
| Faecalibacterium | 0.05398 | 0.1202 |
| Peptostreptococcus | 0 | 0.11515 |
| Porphyromonas | 0.05803 | 0.10303 |
| Capnocytophaga | 0 | 0.09394 |

d) Eigenvector centrality

| Genus | | |
|---|---|---|
| Highest values on the H. lusitanicum group: | H. lusitanicum | R. sanguineus |
| Treponema | 1 | 0.86933 |
| Alistipes | 0.97764 | 0.85859 |
| Porphyromonas | 0.97491 | 0.91576 |
| Filifactor | 0.97201 | 0.78418 |
| Faecalibacterium | 0.95866 | 0.82732 |
| Highest values on the R. sanguineus group: | H. lusitanicum | R. sanguineus |
| Akkermansia | 0.56225 | 1 |
| Campylobacter | 0.70082 | 0.95172 |
| Bacteroides | 0.85395 | 0.95 |
| Streptococcus | 0.93674 | 0.94795 |
| Parabacteroides | 0.082 | 0.94653 |

**Table S8.** Comparison of network properties. Results from testing global network metrics of the networks shown in **Figure 3a** for group differences (via a permutation procedure using 1000 permutations). The computed measures for *H. lusitanicum* and *R. sanguineus* in locality 3, the absolute difference, and the p-value for testing the null hypothesis are shown, respectively, as H 0: |diff| = 0. Global network properties are defined for the whole network an offer an insight into the overall network structure (Peschet et al., 2020).

| **Whole network** | | | | |
|---|---|---|---|---|
| | *H. lusitanicum* | *R. sanguineus* | abs. difference | p-value |
| Number of components | 6.000 | 1.000 | 5.000 | 0.059524 · |
| Clustering coefficient | 0.699 | 0.818 | 0.119 | 0.428571 |
| Modularity | 0.117 | 0.051 | 0.065 | 0.654762 |
| Positive edge percentage | 64.804 | 56.579 | 8.226 | 0.214286 |
| Edge density | 0.336 | 0.668 | 0.331 | 0.261905 |
| Natural connectivity | 0.238 | 0.460 | 0.222 | 0.226190 |

p-values: one-tailed test with null hypothesis diff=0

**Table S9**. Results from testing centrality measures of the networks in Figure 3b for group differences (via a permutation procedure using 1000 permutations). Shown are, respectively, the computed measures for *H. lusitanicum* and *R. sanguineus* in locality 3, the absolute difference, and the p-value for testing the null hypothesis H 0 : |diff| = 0 (Peschet et al., 2020). The p-values are adjusted for multiple testing using the adaptive Benjamini-Hochberg method (Benjamini and Hochberg, 2000), where the proportion of true H 0 is determined according to Langaas et al. (Langaas et al., 2005). These results contain 10 subjects with the highest absolute group difference. All measures are normalized to [0,1].

| | *H. lusitanicum* | *R. sanguineus* | abs. difference | p-value |
|---|---|---|---|---|
| **Degree** (normalized) | | | | |
| *Capnocytophaga* | 0.000 | 0.844 | 0.844 | 0.074453· |
| *Parabacteroides* | 0.111 | 0.778 | 0.667 | 0.083478· |
| *Pyramidobacter* | 0.222 | 0.844 | 0.622 | 0.083478· |
| *Arenimonas* | 0.244 | 0.844 | 0.600 | 0.074453· |
| *Peptostreptococcus* | 0.222 | 0.800 | 0.578 | 0.083478· |
| *Flexilinea* | 0.156 | 0.644 | 0.489 | 0.083478· |
| *Lactobacillus* | 0.333 | 0.778 | 0.444 | 0.083478· |
| *Phocaeicola* | 0.022 | 0.467 | 0.444 | 0.083478· |
| *Lysobacter* | 0.333 | 0.778 | 0.444 | 0.083478· |
| *Coxiella* | 0.000 | 0.444 | 0.444 | 0.083478· |
| **Betweenness centrality** (normalized) | | | | |
| *Dialister* | 0.004 | 0.153 | 0.148 | 1 |
| *Peptostreptococcus* | 0.000 | 0.115 | 0.115 | 1 |
| *Capnocytophaga* | 0.000 | 0.094 | 0.094 | 1 |
| *Leptotrichia* | 0.000 | 0.085 | 0.085 | 1 |
| *Faecalibacterium* | 0.054 | 0.120 | 0.066 | 1 |
| *Anaeroglobus* | 0.000 | 0.056 | 0.056 | 1 |
| *Lysobacter* | 0.009 | 0.061 | 0.051 | 1 |
| *Fretibacterium* | 0.013 | 0.060 | 0.046 | 1 |
| *Porphyromonas* | 0.058 | 0.103 | 0.045 | 1 |
| *Parabacteroides* | 0.000 | 0.040 | 0.040 | 1 |
| **Closeness centrality** (normalized) | | | | |
| Candidatus *Xiphinematobacter* | 1.346 | 11.757 | 10.411 | 0.000000*** |
| *Pseudolabrys* | 1.811 | 11.749 | 9.938 | 0.000000*** |
| *Capnocytophaga* | 0.000 | 7.756 | 7.756 | 0.146255 |
| *Bacteroides* | 2.341 | 7.783 | 5.442 | 0.188043 |
| *Peptostreptococcus* | 1.400 | 6.833 | 5.433 | 0.175507 |
| *Lactobacillus* | 1.690 | 6.728 | 5.038 | 0.233095 |
| *Streptococcus* | 2.709 | 7.340 | 4.630 | 0.233095 |
| *Porphyromonas* | 2.304 | 6.732 | 4.428 | 0.182819 |
| *Dialister* | 1.866 | 6.107 | 4.240 | 0.233095 |
| *Tannerella* | 2.118 | 6.142 | 4.024 | 0.233095 |
| Significance codes:∗∗∗ : 0.001, ∗∗: 0.01, ∗:0.05, .: 0.1 | | | | |

**Table S10.** Jaccard index values express the similarity of the sets of most central nodes and the sets of hub taxa between the two networks to assess how different the two sets of most central nodes are between the groups. Jaccard's index is 0 if the sets are completely different and 1 for exactly equal sets. $P(J \leq j)$ is the probability that Jaccard's index takes for the present total number of taxa a value less than or equal to the calculated index j ($P(J \geq j)$ is defined analogously) (Peschel et al., 2020).

| | Jacc | P(J ≤ Jacc) | P(J ≥ Jacc) |
|---|---|---|---|
| **degree** | 0.222 | 0.231072 | 0.898335 |
| **betweenness centr.** | 0.278 | 0.412243 | 0.768928 |
| **closeness centr.** | 0.333 | 0.608510 | 0.587757 |
| **eigenvec. centr.** | 0.333 | 0.608510 | 0.587757 |
| **hub taxa** | 0.000 | 0.087791 . | 1.000000 |
| Significance codes:∗∗∗ : 0.001, ∗∗: 0.01, ∗:0.05, .: 0.1 | | | |

**Table S11.** Adjusted Rand index assessed similarity between clusterings and range between -1 and 1, where 1 corresponds to identical clustering and 0 to the expected value for two random clusterings. Consequently, positive values imply that two clusters are more similar, and negative values are less similar than expected at random (Peschel et al., 2020).

| **Adjusted Rand index** | |
|---|---|
| | Whole network |
| **ARI** | 0.659 |
| **p-value** | 0.000 |

ARI in [-1,1] with ARI=1: perfect agreement between clusterings; ARI=0: expected for two random clusterings
p-value: permutation test (n=1000) with null hypothesis ARI=0

### Additional results for whole network properties of *H. lusitanicum* and *R. sanguineus* collected at locality 4

**Table S12.** Properties of the networks shown in Figure 3c: a) Frequency table of clusters in the tick species *H. lusitanicum* network in locality 4. b) Frequency table of clusters in the tick species *R. sanguineus* network in locality 4. c) Detected hub nodes in both groups. d)-f) Centrality values of the genera with the highest centrality in decreasing order. The upper part of the table contains the five genera with the highest centrality in *H. lusitanicum* and the lower part with the highest centrality in *R. sanguineus* respectively. Thus, a genus can occur twice in the same table (Peschet el at., 2020).

a) Cluster at *H. lusitanicum*

| Name | Frequency |
|---|---|
| 1 | 25 |
| 2 | 9 |

b) Cluster at *R. sanguineus*

| Name | Frequency |
|---|---|
| 1 | 16 |
| 2 | 13 |
| 3 | 5 |

c) Hub nodes, in alphabetical order. Based on empirical quantiles of centralities.

| *H. lusitanicum* | *R. sanguineus* |
|---|---|
| *Fusobacterium* | *Clostridium ss 1* |
| *Porphyromonas* | *Fusobacterium* |

d) Degree

| Genus | | |
|---|---|---|
| Highest values on the *H. lusitanicum* group: | *H. lusitanicum* | *R. sanguineus* |
| *Prevotella* | 0.87879 | 0.78788 |
| *Filifactor* | 0.87879 | 0.9697 |
| *Udaeobacter* | 0.87879 | 0.90909 |
| *Fusobacterium* | 0.84848 | 1 |
| *Fretibacterium* | 0.84848 | 1 |
| Highest values on the *R. sanguineus* group: | *H. lusitanicum* | *R. sanguineus* |
| *Fusobacterium* | 0.84848 | 1 |
| *Porphyromonas* | 0.81818 | 1 |
| *Fretibacterium* | 0.84848 | 1 |
| *Streptococcus* | 0.75758 | 1 |
| *Dialister* | 0.81818 | 1 |

e) Betweenness centrality

| Genus | | |
|---|---|---|
| Highest values on the *H. lusitanicum* group: | *H. lusitanicum* | *R. sanguineus* |
| Bacteroides | 0.2178 | 0 |
| Clostridium ss 1 | 0.19886 | 0.14962 |
| Gaiella | 0.18182 | 0 |
| Fusobacterium | 0.10795 | 0.00189 |
| Lachnospiraceae NK4A136 | 0.05871 | 0 |
| Highest values on the *R. sanguineus* group: | *H. lusitanicum* | *R. sanguineus* |
| Lactobacillus | 0 | 0.1572 |
| Clostridium ss 1 | 0.19886 | 0.14962 |
| Porphyromonas | 0 | 0.10227 |
| Treponema | 0.02273 | 0.03598 |
| Filifactor | 0.01326 | 0.03409 |

f) Eigenvector centrality

| Genus | | |
|---|---|---|
| Highest values on the *H. lusitanicum* group: | *H. lusitanicum* | *R. sanguineus* |
| *Fusobacterium* | 1 | 0.94597 |
| *Porphyromonas* | 0.9423 | 0.94139 |
| *Filifactor* | 0.93599 | 0.83823 |
| *Pyradomibacter* | 0.93197 | 0.69206 |
| *Prevotella* | 0.93131 | 0.79639 |
| Highest values on the *R. sanguineus* group: | *H. lusitanicum* | *R. sanguineus* |
| *Clostridium ss 1* | 0.76443 | 1 |
| *Fusobacterium* | 1 | 0.94597 |
| *Porphyromonas* | 0.9423 | 0.94139 |
| *Lactobacillus* | 0.6087 | 0.93555 |
| *Prevotella* | 0.90418 | 0.93326 |

**Table S13.** Comparison of network properties. Results from testing global network metrics of the networks shown in **Figure 3a** for group differences (via a permutation procedure using 1000 permutations). The computed measures for *H. lusitanicum* and *R. sanguineus* in locality 4, the absolute difference, and the p-value for testing the null hypothesis are shown, respectively, as H 0: |diff| = 0. Global network properties are defined for the whole network an offer an insight into the overall network structure (Peschet el at., 2020).

| **Whole network** | | | | |
|---|---|---|---|---|
| | *H. lusitanicum* | *R. sanguineus* | abs. difference | p-value |
| Number of components | 1.000 | 1.000 | 0.000 | 1.000000 |
| Clustering coefficient | 0.857 | 9.921 | 0.064 | 0.396040 |
| Modularity | 0.023 | 0.015 | 0.007 | 0.801980 |
| Positive edge percentage | 65.926 | 73.819 | 7.893 | 0.049505* |
| Edge density | 0.684 | 0.891 | 0.207 | 0.207921 |
| Natural connectivity | 0.416 | 0.479 | 0.063 | 0.495050 |

p-values: one-tailed test with null hypothesis diff=0

**Table S14.** Results from testing centrality measures of the networks in Figure 3b for group differences (via a permutation procedure using 1000 permutations). Shown are, respectively, the computed measures for *H. lusitanicum* and *R. sanguineus* in locality 4, the absolute difference, and the p-value for testing the null hypothesis H 0 : |diff| = 0 (Peschet et al., 2020). The p-values are adjusted for multiple testing using the adaptive Benjamini-Hochberg method (Benjamini and Hochberg, 2000), where the proportion of true H 0 is determined according to Langaas et al. (2005). These results contain 10 subjects with the highest absolute group difference. All measures are normalized to [0,1].

| | *H. lusitanicum* | *R. sanguineus* | abs. difference | p-value |
|---|---|---|---|---|
| **Degree** (normalized) | | | | |
| *Peptostreptococcus* | 0.061 | 0.970 | 0.909 | 0.264339 |
| *Francisella* | 0.030 | 0.727 | 0.697 | 0.634413 |
| *Alistipes* | 0.121 | 0.818 | 0.697 | 0.490915 |
| *Johnsonella* | 0.061 | 0.727 | 0.667 | 0.634413 |
| *Lachnospiraceae NK4A136 group* | 0.333 | 0.939 | 0.606 | 0.490915 |
| *Bacteroides* | 0.333 | 0.848 | 0.515 | 0.490915 |
| *Lactobacillus* | 0.667 | 0.970 | 0.303 | 0.490915 |
| *Streptococcus* | 0.758 | 1.000 | 0.242 | 0.490915 |
| *Clostridium sensu stricto 1* | 0.758 | 0.970 | 0.212 | 0.594762 |
| *Coxiella* | 0.818 | 0.606 | 0.212 | 0.674520 |
| **Betweenness centrality** (normalized) | | | | |
| *Bacteroides* | 0.218 | 0.000 | 0.218 | 0.104377 |
| *Gaiella* | 0.182 | 0.000 | 0.182 | 0.104377 |
| *Lactobacillus* | 0.000 | 0.157 | 0.157 | 0.104377 |
| *Fusobacterium* | 0.108 | 0.002 | 0.106 | 0.626264 |
| *Porphyromonas* | 0.000 | 0.102 | 0.102 | 0.626264 |
| *Lachnospiraceae NK4A136 group* | 0.059 | 0.000 | 0.059 | 0.313132 |
| *Capnocytophaga* | 0.051 | 0.000 | 0.051 | 0.626264 |
| *Clostridium sensu stricto 1* | 0.199 | 0.150 | 0.049 | 0.626264 |
| *Peptostreptococcus* | 0.000 | 0.027 | 0.027 | 0.626264 |
| *Prevotella* | 0.000 | 0.021 | 0.021 | 0.917029 |
| **Closeness centrality** (normalized) | | | | |
| *Clostridium sensu stricto 1* | 1.706 | 3.774 | 2.068 | 0.666668 |
| *Lactobacillus* | 1.281 | 3.011 | 1.730 | 0.333334 |
| *Fusobacterium* | 2.895 | 1.472 | 1.423 | 0.968752 |
| *Porphyromonas* | 2.360 | 3.775 | 1.414 | 0.968752 |
| *Tannerella* | 2.421 | 1.437 | 0.983 | 0.968752 |
| *Flexilinea* | 1.956 | 1.128 | 0.828 | 0.968752 |
| *Fretibacterium* | 2.288 | 1.476 | 0.812 | 0.968752 |
| *Anaeroglobus* | 1.914 | 1.177 | 0.737 | 0.968752 |
| *Francisella* | 0.000 | 0.665 | 0.665 | 0.968752 |
| *Johnsonella* | 0.365 | 0.926 | 0.561 | 0.968752 |
| Significance codes:∗∗∗ : 0.001, ∗∗: 0.01, ∗:0.05, .: 0.1 | | | | |

**Table S15.** Jaccard index values express the similarity of the sets of most central nodes and the sets of hub taxa between the two networks to assess how different the two sets of most central nodes are between the groups. Jaccard's index is 0 if the sets are completely different and 1 for exactly equal sets. P(J ≤ j) is the probability that Jaccard's index takes for the present total number of taxa a value less than or equal to the calculated index j (P(J ≥ j) is defined analogously) (Peschel et al., 2020).

| | Jacc | P(J ≤ Jacc) | P(J ≥ Jacc) |
|---|---|---|---|
| **degree** | 0.222 | 0.377178 | 0.856932 |
| **betweenness centr.** | 0.286 | 0.475500 | 0.738807 |
| **closeness centr.** | 0.385 | 0.758692 | 0.447961 |
| **eigenvec. centr.** | 0.636 | 0.991177 | 0.038629 * |
| **hub taxa** | 0.333 | 0.740741 | 0.703704 |
| Significance codes:∗∗∗ : 0.001, ∗∗: 0.01, ∗:0.05, .: 0.1 | | | |

**Table S16.** Adjusted Rand index assessed similarity between clusterings and range between -1 and 1, where 1 corresponds to identical clustering and 0 to the expected value for two random clusterings. Consequently, positive values imply that two clusters are more similar, and negative values are less similar than expected at random (Peschel et al., 2020).

| Adjusted Rand index | |
|---|---|
| | **Whole network** |
| **ARI** | 0.220 |
| **p-value** | 0.003 |

ARI in [-1,1] with ARI=1: perfect agreement between clusterings; ARI=0: expected for two random clusterings
p-value: permutation test (n=1000) with null hypothesis ARI=0
